# IFN-γ-Dependent Macrophage Reprogramming Coordinates Inflammatory Resolution and Matrix Remodeling in Heart Regeneration

**DOI:** 10.64898/2026.03.18.712551

**Authors:** Khai Lone Lim, Kaushik Chowdhury, Yu-Jen Hung, Ben Shih-Lei Lai

## Abstract

Heart regeneration requires coordinated immune activation, timely inflammatory resolution, and dynamic extracellular matrix (ECM) remodeling in addition to cardiomyocyte (CM) proliferation. However, the cytokine signals that instruct immune cell functions during cardiac repair remain incompletely understood. Here, we identify interferon-gamma (IFN-γ) as a critical regulator of macrophage plasticity in zebrafish heart regeneration. IFN-γ signaling components are dynamically activated following cardiac injury, with early induction of *ifng1* and temporally coordinated receptor expression. Genetic ablation of *ifng1* impairs myocardial regeneration, resulting in reduced CM proliferation and persistent fibrotic scarring. Temporal transcriptional profiling reveals sustained inflammatory signatures, impaired efferocytosis, and abolished reparative programs, accompanied by aberrant immune cell dynamics and retention of injury-derived debris in mutant hearts. Transcriptomic analysis of cardiac macrophages further reveals that IFN-γ deficiency disrupts the transition from an inflammatory state to a reparative, ECM-remodeling phenotype, leading to reduced collagen denaturation and diminished CM protrusion at the injury border zone. Inducible- and macrophage-specific blockade of IFN-γ signaling phenocopies defects in global knockout, establishing a cell-autonomous requirement for IFN-γ in coordinating regenerative immune function. Collectively, our findings define an IFN-γ-dependent macrophage reprogramming axis that couples inflammatory resolution to ECM remodeling in heart regeneration, elucidating how cytokine signaling actively instructs tissue repair.

**GRAPHICAL ABSTRACT:** 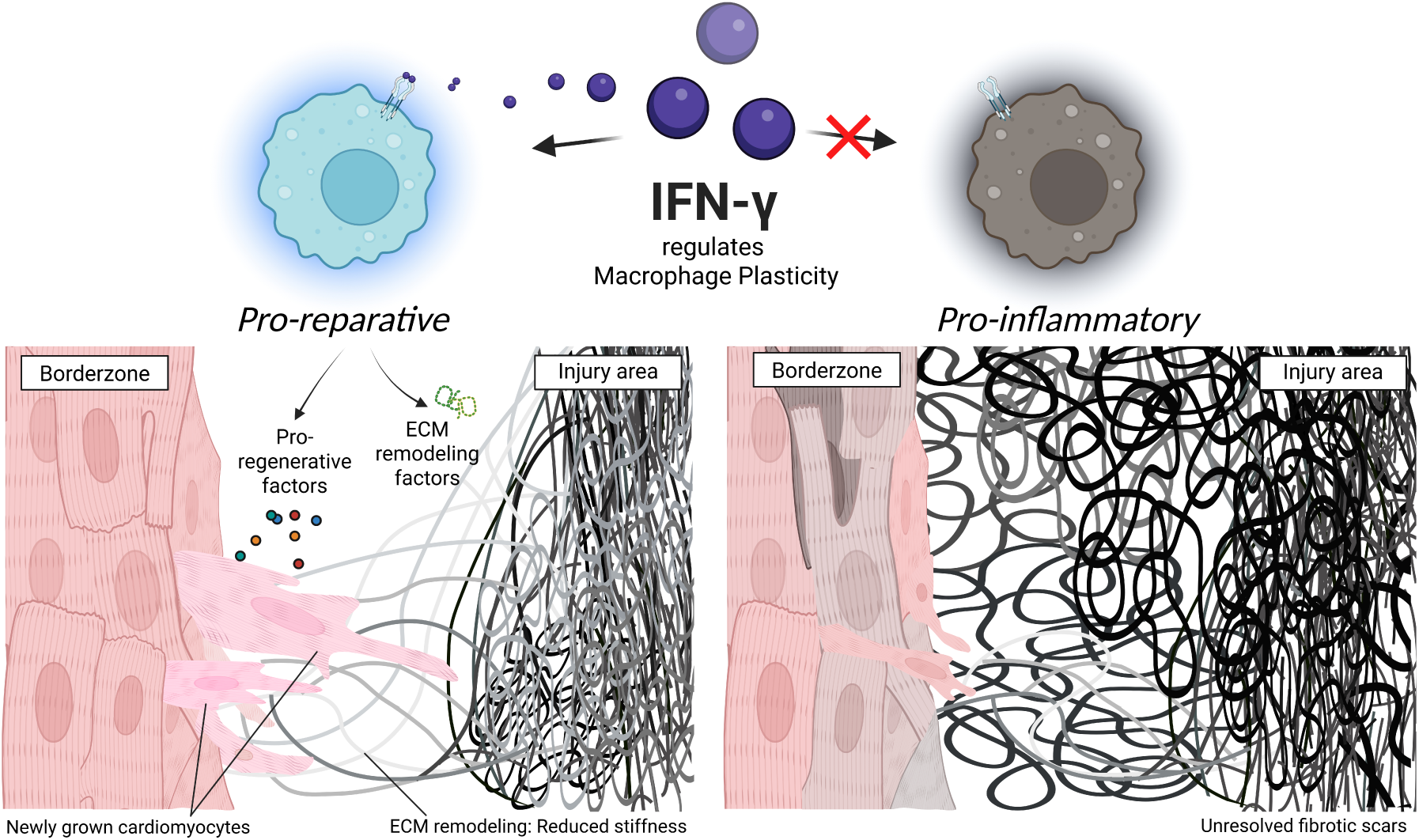

## INTRODUCTION

Heart failure (HF) affects more than 64 million people worldwide and remains a leading cause of morbidity and mortality^1^. Over 30% of HF cases arise from atherothrombotic myocardial infarction (MI)^2^. Although advances in acute and post-MI care have markedly reduced early mortality^3,4^, most MI survivors ultimately develop HF, reflecting the severely limited regenerative capacity of the adult mammalian heart^5,6^. Rather than regenerating lost myocardium, the adult mammalian heart responds to myocardial infarction with fibrotic scar formation, leading to adverse remodeling, progressive contractile dysfunction, and eventual heart failure.

In contrast, certain vertebrates, including some fish and amphibians, retain an intrinsic capacity for heart regeneration after comparable injury^7^. Among these, zebrafish (*Danio Rerio*) exhibit a remarkable regenerative response, activating cardiomyocyte (CM) proliferation and ultimately replacing scar tissue to fully restore cardiac structure and function after injury^8–10^. Notably, this regenerative capacity is absent in another teleost fish, the medaka (*Oryzias latipes*), despite similar cardiac anatomy, physiology, environmental conditions, and close phylogenetic relationship^11–14^. To dissect the cellular and molecular basis of heart regeneration, we established a comparative platform using regenerative zebrafish and non-regenerative medaka. Our previous studies revealed striking differences in immune responses and coronary vessel formation between these species and identified macrophage dynamics and function as central determinants of regenerative outcome^12^. Importantly, modulating immune response using poly(I:C), a pharmaceutical immunostimulant identified through zebrafish-medaka comparison, was sufficient to induce *de novo* heart regeneration in medaka, in a macrophage-dependent manner, highlighting macrophages as both essential and instructive regulators of cardiac regeneration^12^.

The requirement for macrophages in heart regeneration is evolutionarily conserved beyond teleosts. In neonatal mice, macrophage depletion also impairs regenerative capacity following cardiac injury^15^. Neonatal macrophages distinctly respond to stressed CMs and secrete mitogenic factors that stimulate CM proliferation, underscoring a critical cardiac-immune crosstalk^16^. Consistent with this concept, our recent studies identified specialized subsets of cardiac resident macrophages that coordinate inflammation resolution, debris clearance, ECM remodeling, and CM replenishment during zebrafish heart regeneration^17^. Furthermore, we demonstrated that poly(I:C) administration promotes heart regeneration in medaka via macrophage-dependent mechanisms (Chowdhury et al., 2026, *manuscript in revision*). Together, these observations position macrophage function as a conserved and central axis governing cardiac repair and regeneration.

While poly(I:C) effectively stimulates a regenerative response, it remains an exogenous synthetic immunostimulant. In contrast, interferon-gamma (IFN-γ) emerged from our comparative analysis as a key endogenous upstream regulator preferentially enriched in zebrafish over medaka following cardiac injury^12^. Notably, IFN-γ signaling shares common downstream effectors with the poly(I:C)-induced type I IFN response, positioning it as a physiological driver of cardiac regeneration^18,19^. IFN-γ is a type II interferon primarily produced by T cells, natural killer (NK) cells, and, in certain contexts, macrophages, when stimulated by cytokines such as IL-12, IL-15, and IL-18, as well as by type I interferons downstream of poly(I:C) stimulation^18,20,21^. IFN-γ is conventionally recognized for its role in antiviral defense and immune regulation, particularly through macrophage activation, efferocytosis, and major histocompatibility complex class II expression^22–24^. IFN-γ has been implicated in cardiac repair after MI, with elevated circulating levels associated with poorer repair outcomes^25^. In a zebrafish and a mouse model of ion imbalance and pressure overload, IFN-γ has also been implicated to drive macrophage reprogramming, cerebrovascular remodeling, and cognitive dysfunction^26^. Paradoxically, IFN-γ–deficient mice display extensive scarring and impaired functional recovery after MI, accompanied by reduced myeloid cell activation and infiltration, suggesting a beneficial role for IFN-γ in cardiac repair^27^. These seemingly opposing effects highlight a critical gap in understanding how IFN-γ signaling is dynamically regulated and whether it contributes to optimal cardiac repair and regeneration remains poorly characterized. Here, we hypothesized that IFN-γ plays an important role in zebrafish heart regeneration and systematically characterized its activation, functional requirement, and cellular and molecular mechanisms, identifying how IFN-γ coordinates macrophage function in heart regeneration.

## RESULTS

### Dynamic activation of IFN-γ following cardiac injury is functionally essential for heart regeneration

To assess the involvement of IFN-γ signaling during zebrafish heart regeneration, we first examined the temporal expression of the IFN-γ ligand (*ifng1*) and its receptors (*crfb6*, *crfb13*, *crfb17*)^28^ following cardiac injury. Quantitative reverse-transcription PCR (RT-qPCR) revealed a rapid activation of *ifng1* as early as 6-hour post-cardiac injury (hpci), with a pronounced peak at 1-day post-cardiac injury (dpci) (Fig. 1A). After returning to basal level between 3-7 dpci, a second activation of *ifng1* was observed at 14 dpci. Interestingly, the activation of the IFN-γ receptor genes^28^, *crfb6* (*ifngr2*), *crfb13* (*ifngr1l*) and *crfb17* (*ifngr1*) corresponded closely with the dynamics of *ifng1* expression (Fig. 1A). Consistent with these findings, *in situ* hybridization confirmed the peak expression of *ifng1* at 1 dpci, which predominantly localized to the injured area (Fig. 1B).

**Fig. 1.**
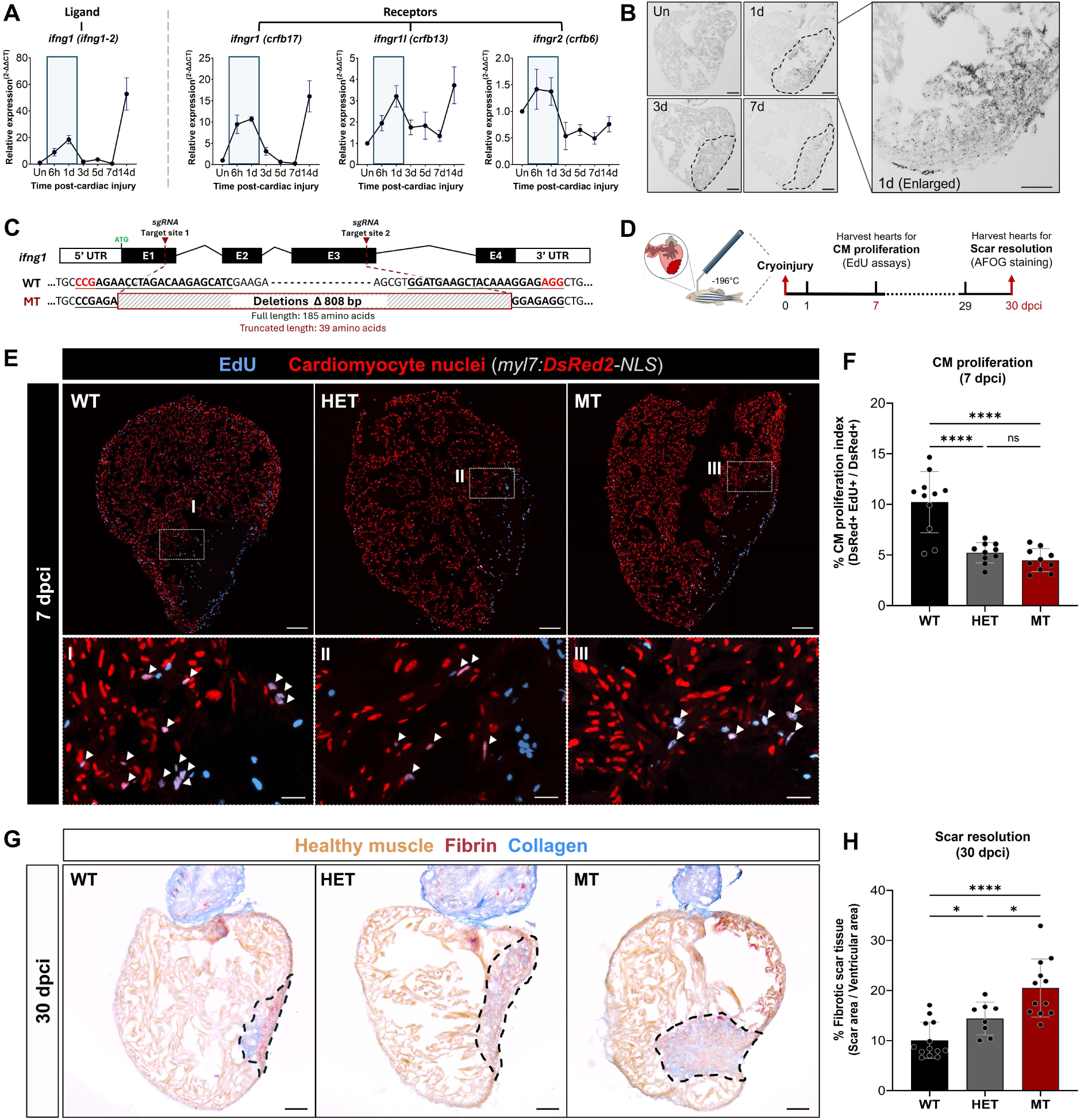
Reactivation of *ifng1* is required for zebrafish heart regeneration. **A** Quantitative PCR analysis of IFN-γ ligand and receptor expression in whole zebrafish ventricles at untouched (Un), 6 h, 1, 3, 5, and 7 days post-cardiac injury (dpci) (n = 3 ventricles per group). **B** In situ hybridization showing *ifng1* expression in zebrafish hearts at Un, 1, 3 and 7 dpci. Higher-magnification image at 1 dpci shows *ifng1* expression predominantly localized to the injury region and extending across the border zone. Scale bar: 100 μm. **C** Generation of the *ifng1* knockout zebrafish line using CRISPR–Cas9–mediated mutagenesis. Schematic of the *ifng1* genomic locus containing four exons. Red arrowheads denote single-guide RNA (sgRNA) target sites in exons 1 and 3. All sgRNA target sequences are shown in bold and underlined (uppercase), with PAM sequences highlighted in red. The resulting large genomic deletion introduces a premature stop codon, leading to early truncation of the IFNγ protein (full length: 189 amino acids; truncated length: 39 amino acids). **D** Experimental design for assessing cardiac regeneration following cardiac cryoinjury. Hearts were collected at 7 dpci to evaluate cardiomyocyte (CM) proliferation and at 30 dpci to assess scar resolution. **E** Representative heart sections from wild-type (WT), heterozygous (*ifng1*^+/−^; HET) and mutant (*ifng1*^−/−^; MT) hearts at 7 dpci, stained for proliferating cells (light blue) and CM nuclei (red). Proliferating CMs are indicated by overlapping signals (violet). Scale bars: 100 μm (overview) and 20 μm (insets). **F** Quantification of CM proliferative index (%) in the border zone of WT (n = 11), HET (n = 10) and MT (n = 10) hearts at 7 dpci. For each heart, up to three sections containing the largest areas were analyzed. **G** Representative AFOG-stained heart sections at 30 dpci. Myocardium is shown in orange–brown, fibrin in red and collagen in blue. Scar areas were outlined by dashed lines. Scale bar: 100 μm. **H** Quantification of fibrotic scar area (%) in WT (n = 13), HET (n = 8) and MT (n = 12) hearts at 30 dpci. For each heart, up to three sections containing the largest areas were analyzed. Data in **F** and **H** are presented as mean ± S.D. Statistical significance was determined using one-way ANOVA followed by Tukey’s multiple-comparison test (\**P* < 0.05, \*\*\*\**P* < 0.0001; ns, not significant).

We performed a meta-analysis of published bulk^12^ and single-cell RNA-sequencing datasets^14,17,29^ and observed similar activation of *ifng1* and its receptors (*crfb6*, *crfb13* and *crfb17*) following injury (Supplementary Fig. 1B). Cell-type–resolved analysis of scRNA-seq data further revealed that *ifng1* expression was predominantly enriched in T-cell populations, with some expression in resident macrophage subsets (Supplementary Fig. 1C,D), while the receptors are more broadly expressed in macrophages and other cardiac cells (Supplementary Fig.1C,D). Interestingly, we also observed activation of *ifng1* in medaka following regenerative poly (I:C) stimulation. In this context, *ifng1* was likewise induced predominantly in T-cells and macrophage subsets (Supplementary Fig. 1E; (Chowdhury et al., 2026, *manuscript in revision*). These findings suggest that core components of IFN-γ signaling are dynamically activated following cardiac injury, associated with regenerative capacity across species.

To determine the functional requirement of IFN-γ signaling in zebrafish heart regeneration, we generated an *ifng1* knockout line using CRISPR/Cas9-mediated genome editing (Fig. 1C). Homozygous mutant (MT) derived from heterozygous (HET) incrosses developed into adulthood without overt morphological abnormalities, and their frequency conformed to the expected Mendelian ratio (Supplementary Fig. 2). To determine the impact of IFN-γ deficiency on cardiac regeneration, we performed cryoinjury and quantified CM proliferation at 7-day post cardiac injury (dpci) and fibrotic scar size at 30 dpci. Compared with wild-type WT siblings, both HET and MT hearts exhibited a significantly reduced CM proliferation (Fig. 1E, quantified in F). Correspondingly, both HET and MT hearts exhibited significantly larger fibrotic scars than WT controls at 30 dpci (Fig. 1G, quantified in H). These results suggest that IFN-γ signaling is required for efficient cardiomyocyte proliferation and successful heart regeneration following injury.

### Temporal transcriptomic profiling reveals dysregulated repair programs in *ifng1* mutant hearts

To investigate the mechanisms underlying fibrotic repair in *ifng1* MT hearts, we performed bulk RNA sequencing across multiple time points (untouched, 6 h, 1 d, 3 d, 5 d, and 7 dpci) (Fig. 2A). Principal component analysis (PCA) showed that initial transcriptional responses in *ifng1* MT hearts were similar but to a lesser extent compared to WT hearts before 3-5 dpci. Strikingly, from 5 dpci onward, *ifng1* MT hearts diverged from WT trajectories and exhibited premature termination of the reparative program, such that by 7 dpci their transcriptomic profiles closely resembled the uninjured state (Fig. 2B). Hierarchical clustering of differentially expressed genes (DEGs) further revealed sustained and aberrant activation of leukocyte- and immune-associated gene programs in *ifng1* MT hearts at 7 dpci (Fig. 2C, HC1–4), accompanied by diminished induction of reparative pathways, including genes involved in heart morphogenesis and extracellular matrix remodeling (Fig. 2C, HC5–8). Notably, we also detected dysregulated activation of angiogenesis- and endothelial-related gene signatures in *ifng1* MT hearts (Fig. 2C, HC15–16). Together, these findings suggest that IFN-γ signaling exerts pleiotropic effects during cardiac repair, acting directly or indirectly on multiple biological processes involving cardiac cell types, including immune cells, cardiomyocytes, and endothelial cells, and coordinating their responses to enable effective heart regeneration.

**Fig. 2.**
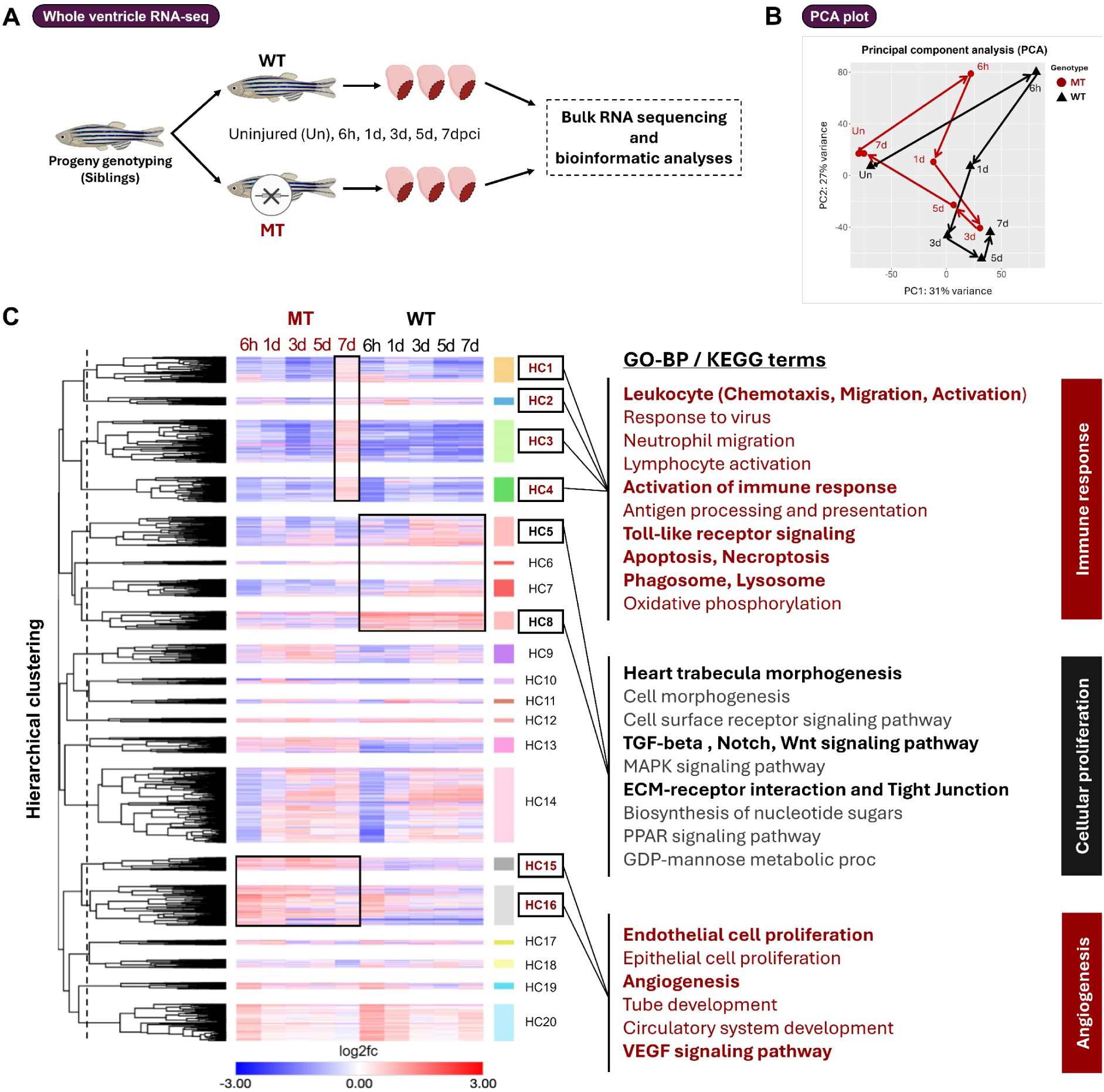
Transcriptomic profiling of WT and *ifng1* MT hearts. **A** Zebrafish ventricles were collected at uninjured (Un), 6 h, 1 d, 3 d, 5 d and 7 dpci and subjected to RNA sequencing (n = 4 ventricles per group). **B** Principal component analysis (PCA) revealed that *ifng1* MT hearts begin to diverge from the WT regenerative transcriptome at 3 dpci and continued to diverge thereafter. **C** Hierarchical clustering differentially expressed genes (DEGs) across time points in WT and *ifng1* MT hearts. GO and KEGG enrichment of upregulated clusters highlight terms associated with immune response, cellular proliferation, and angiogenesis.

### IFN-γ deficiency disrupts immune cell dynamics during cardiac repair

Enlightened by the hint of immune dysregulation from transcriptomic profiling, we next examined immune cellular dynamics in WT and *ifng1* MT hearts (Fig. 3). Using the *Tg(mpx:EGFP);Tg(mpeg1.1:mCherryF)*^30,31^ reporter line, we observed that while initial innate immune cell recruitment at 1-3 dpci was comparable between genotypes, the subsequent resolution phase was significantly impaired in the absence of IFN-γ (Fig. 3A, quantified in C and D). In WT hearts, both macrophages^32^ and neutrophils^33^ infiltrated the injured area after injury, while neutrophils began to resolve by 3 dpci. In contrast, *ifng1* MT hearts exhibited exaggerated macrophage accumulation and neutrophil retention at 7 dpci (Fig. 3C,D). These findings suggest a defect in the transition from inflammatory recruitment to resolution, resulting in sustained inflammatory cell presence at the injury site. Analysis using the *TgBac(lck:EGFP)*^34^ reporter further revealed altered adaptive immune dynamics in *ifng1* MT hearts. While T-cells robustly infiltrate WT hearts between 7 and 14 dpci, similar to previous observations^35,36^, T-cell numbers were significantly reduced in *ifng1* MT hearts (Fig. 3B, quantified in E). Together, dysregulated innate resolution and altered adaptive recruitment align with the transcriptomic analyses and support a role for IFN-γ signaling in coordinating immune cell dynamics and timely inflammatory resolution during cardiac repair.

**Fig. 3.**
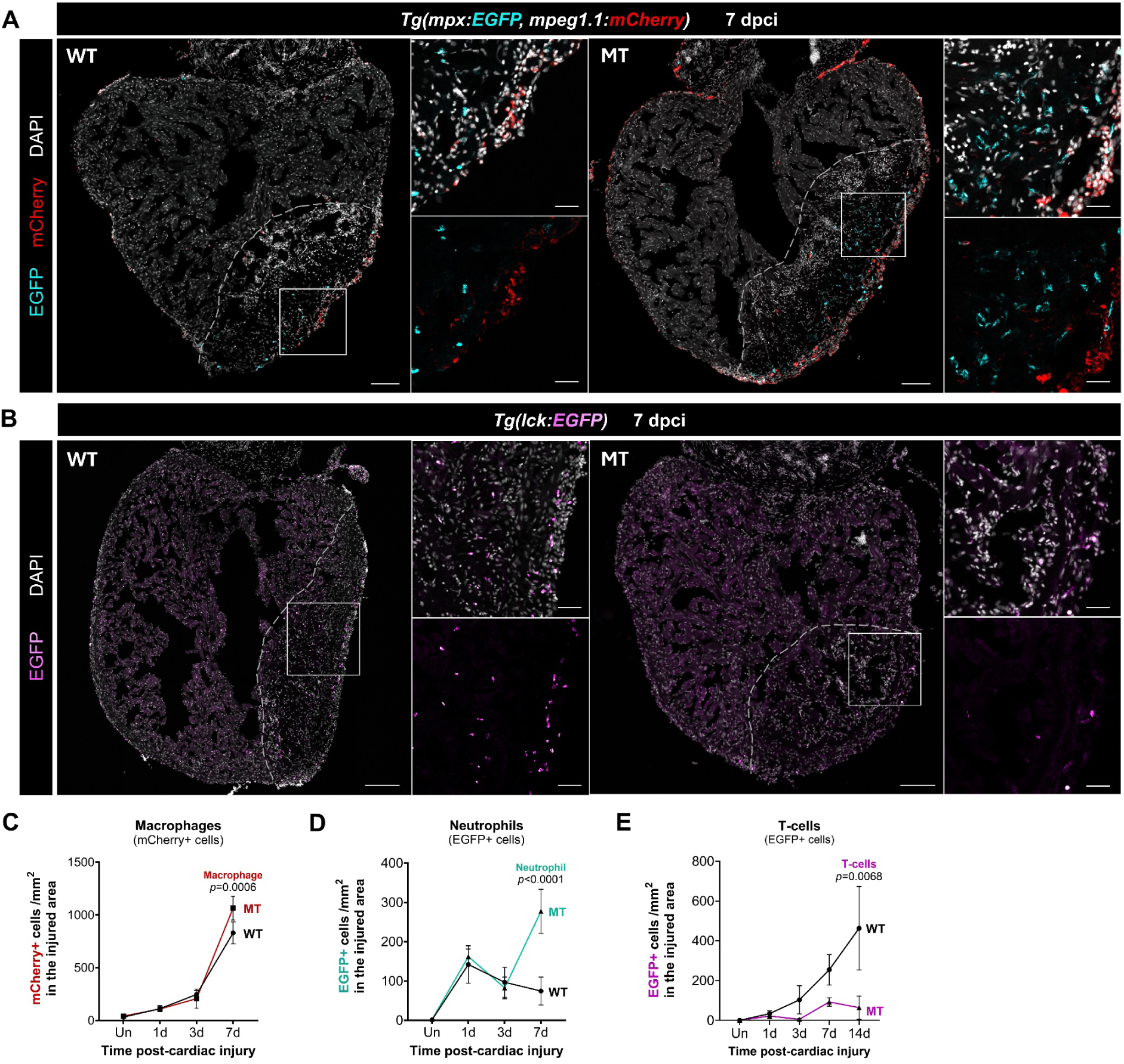
Dynamics of immune cell infiltration in WT and *ifng1* MT hearts following cardiac injury. **A** Representative immunofluoresence images of neutrophils (cyan, *mpx:*EGFP) and macrophages (red, *mpeg1.1:*mCherryF) in WT and *ifng1* MT heart sections at 7 dpci. Scale bars: 100 μm (overview) and 25 μm (insets). Sample sizes per group: Un (n=3), 1 dpci (n=3), 3dpci (n=4), and 7 dpci (n=4). **B** Representative immunofluorescence images of T cells and T-cell precursors (magenta, *lck:*EGFP) in WT and *ifng1* MT heart sections at 7 dpci. Scale bars: 100 μm (overview) and 25 μm (insets). Sample sizes per group: Un (n=3), 1 dpci (n=3), 3dpci (n=4), 7 dpci (n=4) and 14 dpci (n=4). **C-E** Quantification of immune cell recruitment to the injury area across time points, showing densities of **C**, macrophages; **D**, neutrophils; and **E**, T cells. Data are presented as mean ± S.D. Statistical significance was assessed by two-way ANOVA followed by Holm–Šídák multiple-comparisons test.

### IFN-γ signaling is required for macrophage-mediated debris clearance and inflammatory resolution

Macrophages play central roles in debris clearance, inflammatory resolution, and extracellular matrix (ECM) remodeling during cardiac repair and regeneration^12,17,32,35,37–43^. We therefore hypothesized that IFN-γ signaling regulates essential macrophage functions during cardiac repair. To assess macrophage-mediated debris clearance under IFN-γ–deficient conditions, we performed phalloidin staining to visualize filamentous actin (F-actin) within the injury site in *ex vivo*-cultured hearts^44^. In this context, exposed F-actin serves as a marker of injury-derived material and necrotic cellular debris^45^. In uninjured hearts, minimal phalloidin signal was detected in both WT and *ifng1* MT hearts, consistent with the absence of exposed F-actin in intact tissue (Fig. 4A). Following injury, robust phalloidin labeling was observed within the injured region of both genotypes at 1 dpci. In WT hearts, this debris-associated signal was markedly reduced by 3 dpci, reflecting efficient clearance of injury-derived material (Fig. 4A, quantified in B). In contrast, phalloidin-positive debris persisted prominently in *ifng1* MT hearts from 3 to 7 dpci, indicating delayed clearance and impaired resolution of injury-associated material in the absence of IFN-γ signaling (Fig. 4A, quantified in B). Notably, this persistence occurred despite the increased macrophage accumulation observed in *ifng1* MT hearts (Fig. 3C), suggesting that macrophage functional capacity, rather than recruitment, is compromised under IFN-γ-deficient conditions. The delayed removal of injury-derived debris parallels sustained neutrophil retention (Figure 3D) and immune-related transcriptomic signatures in *ifng1* MT hearts (Fig. 2C), collectively supporting a model in which IFN-γ deficiency leads to macrophage dysfunction and defective inflammatory resolution during heart regeneration.

**Fig. 4.**
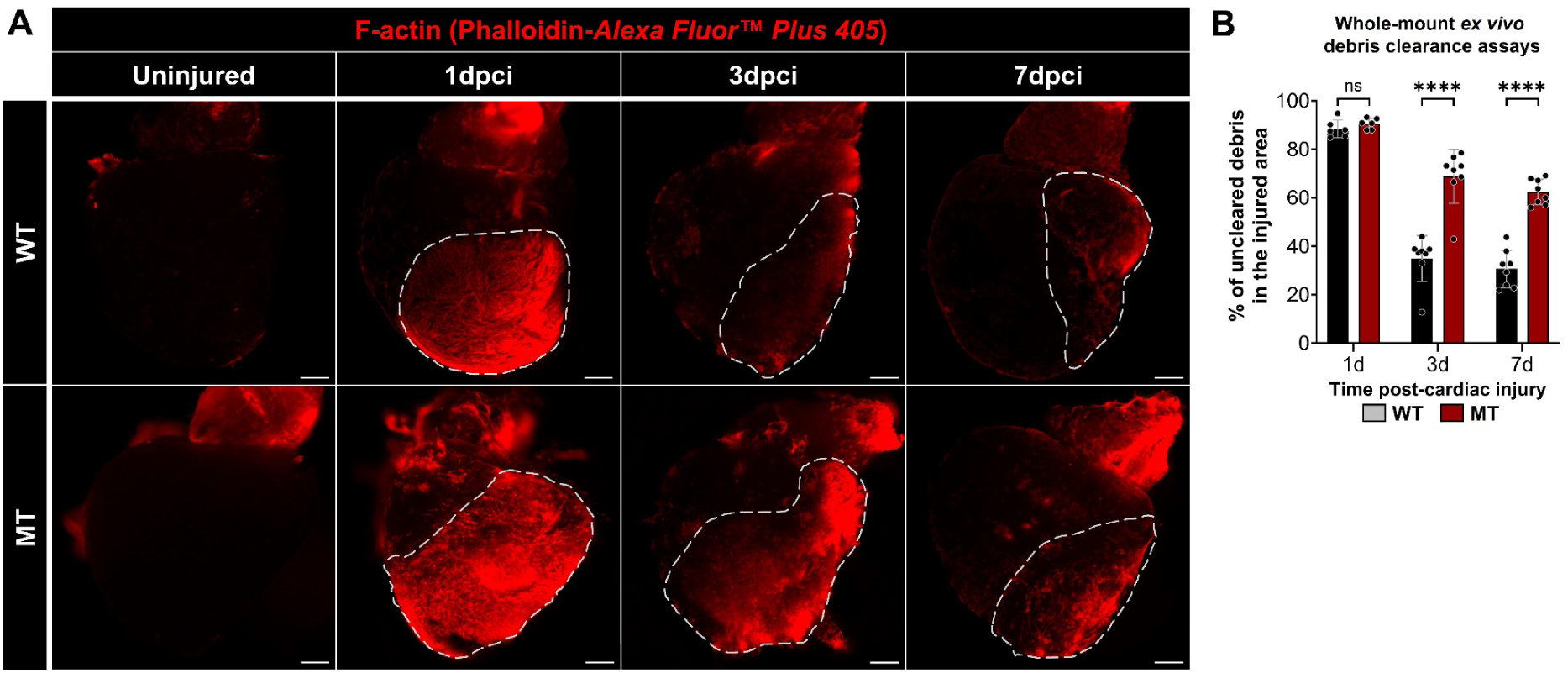
*ifng1* deficiency impairs macrophage-mediated debris clearance in the injured area. **A** Representative whole-mount *ex vivo* images of WT and *ifng1* MT ventricles stained with Alexa Fluor™ Plus 405 Phalloidin (red) to visualize F-actin+ cellular debris within the injury area. Scale bar: 150 µm. **B** Quantification of uncleared debris, expressed as a percentage (%) of the total injured area coverage, at 1, 3, and 7 dpci. Sample sizes: 1 dpci (*n* = 6), 3 dpci (*n* = 8), and 7 dpci (*n* = 8). Data are mean ± SD. Statistical significance was determined by two-way ANOVA with Holm-Šídák’s multiple comparison test. (\*\*\*\**P* < 0.0001; ns, not significant).

### IFN-γ deficiency disrupts Macrophage transcriptional reprogramming during cardiac repair

To investigate the molecular mechanisms by which IFN-γ regulates macrophage function during cardiac repair, we performed RNA sequencing on FACS-isolated cardiac macrophages from WT and *ifng1* MT hearts at 3 and 7 dpci (Fig. 5A). Unbiased hierarchical clustering of normalized transcriptomic profiles revealed a pronounced divergence in macrophage activation states between genotypes (Fig. 5B). In WT macrophages, we observed a marked temporal shift in gene expression between 3 and 7 dpci, corresponding to previously described inflammatory and tissue-repair phases^46–53^. At 3 dpci, WT macrophages upregulated genes associated with injury response and DNA repair (HC1), efferocytosis and metabolic adaptation (HC2)^54,55^, and inflammatory activation (HC3, HC8). By 7 dpci, this program transitioned toward activation of genes involved in pro-reparative signaling (HC4) and extracellular matrix (ECM) remodeling (HC5). In contrast, this phenotypic transition was substantially disrupted in *ifng1* MT macrophages, which showed reduced induction of efferocytosis-associated (HC2), pro-reparative (HC4), and ECM remodeling genes (HC5), alongside sustained inflammatory and neutrophil-activating signatures (HC3, HC8, HC9).

**Fig. 5.**
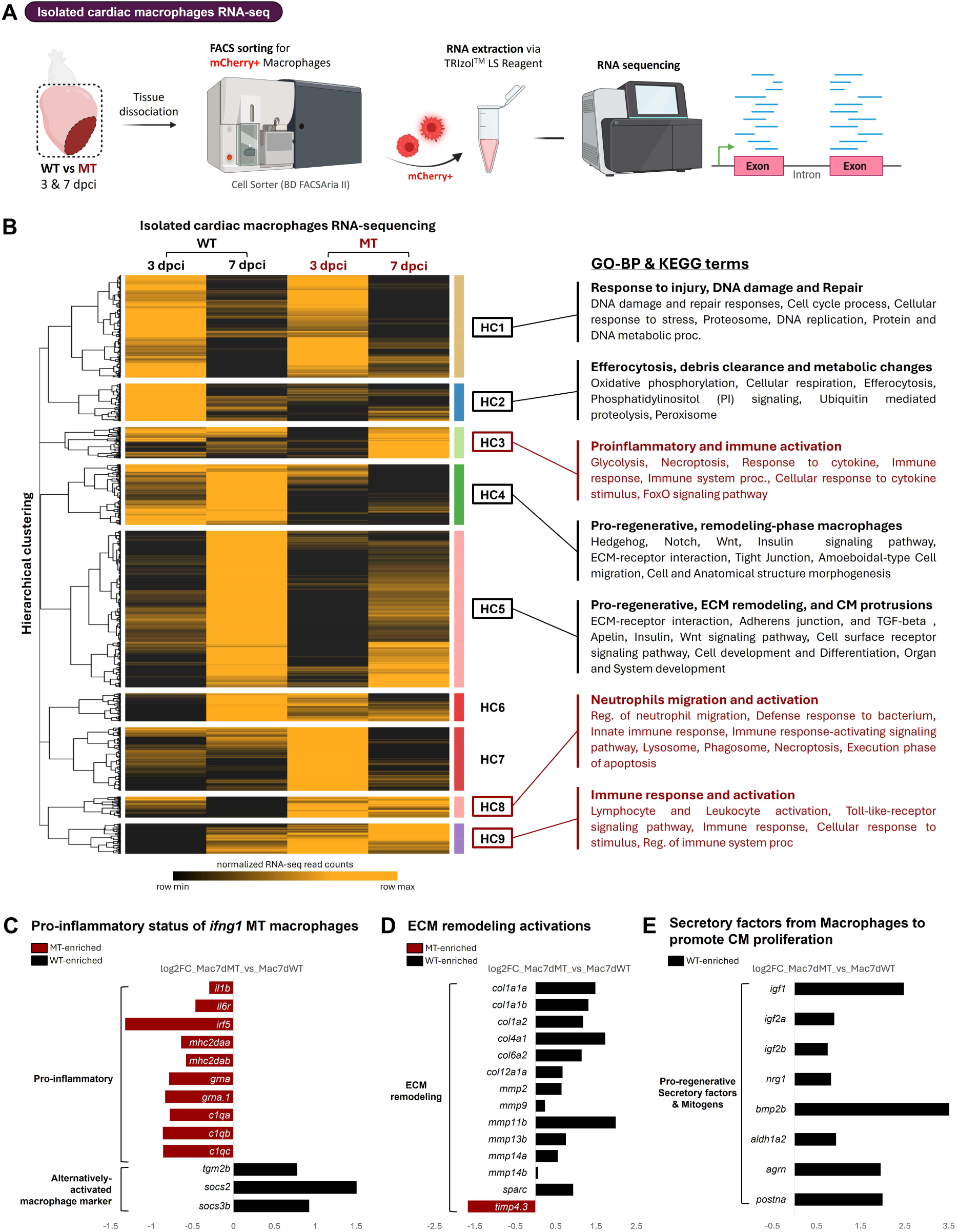
Transcriptomic profiling of isolated cardiac macrophages from WT and *ifng1* MT hearts. **A** Experimental schematic illustrating the isolation of cardiac macrophages from WT and *ifng1* MT hearts at 3 and 7 dpci, followed by fluorescence-activated cell sorting (FACS) and bulk RNA-sequencing. **B** Unbiased hierarchical clustering of normalized transcriptomes from isolated cardiac macrophages. Annotated Gene Ontology-Biological Process (GO-BP) and KEGG pathways highlight enriched terms associated with inflammatory response and tissue remodeling. **C-E** Barplots showing log2 fold-change (Log2FC) of differentially expressed genes associated with: **C,** pro-inflammatory and alternatively-activated macrophage markers; **D,** ECM remodeling; and **E**, Pro-regenerative secretory factors or mitogens.

Differential gene expression analysis at 7 dpci further substantiated these genotype-specific states. Compared with WT macrophages, *ifng1* MT macrophages showed persistent upregulation of pro-inflammatory mediators, including cytokine-related genes (*il1b*, *il6r*, *irf5*), antigen presentation molecules (*mhc2daa*, *mhc2dab*), and complement components (*c1qa*, *c1qb*, *c1qc*)(Fig. 5C), consistent with unresolved inflammatory activation. Conversely, WT macrophages preferentially expressed genes associated with tissue remodeling, including ECM components (*col1a1a*, *col1a1b*, *col1a2*, *col4a1*, *col6a2*, *col12a1a*)^32,56–58^, matrix metalloproteinases (*mmp2, mmp9, mmp11b, mmp14b,* and *mmp15b*)^32,43,59–62^, and the matricellular protein (*sparc*)^56,63^ (Fig. 5D). Notably, WT macrophages also exhibited elevated expression of secreted factors implicated in CM proliferation and regeneration, including *igf1/2, nrg1, bmp2b, aldh1a2, agrn, and postna* (Fig. 5E)^64–74^. Collectively, these findings indicate that IFN-γ signaling is required for the timely transition of macrophages from an inflammatory state to a reparative phenotype that supports cardiac regeneration.

### IFN-γ signaling Is required for dynamic ECM remodeling and CM protrusion at the injury border zone

Dynamic ECM remodeling is a hallmark of successful heart regeneration, characterized by the transition from fibrin deposition to collagen accumulation, followed by collagen degradation and replacement by regenerating CMs^9,59,60^. To functionally assess macrophage-mediated ECM remodeling under IFN-γ–deficient conditions, we examined spatial and temporal dynamics of collagen turnover using collagen-hybridizing peptide (CHP) staining at 7 and 14 dpci in WT and *ifng1* MT hearts (Fig. 6). Consistent with previous observations, fibrin was fully replaced by collagen by 7 dpci in WT hearts, while prominent CHP-positive collagen degradation/remodeling was detected at the injury border zone, a region previously shown to be enriched for resident macrophages (Fig. 6A)^43^. This remodeling activity progressively expanded to encompass the entire injured area by 14 dpci (Fig. 6B,C, quantified in D), reflecting coordinated scar remodeling during regeneration. In contrast, *ifng1* MT hearts exhibited markedly reduced CHP staining at both 7 and 14 dpci (Fig. 6A–C, quantified in D), indicating impaired collagen degradation and remodeling.

**Fig. 6.**
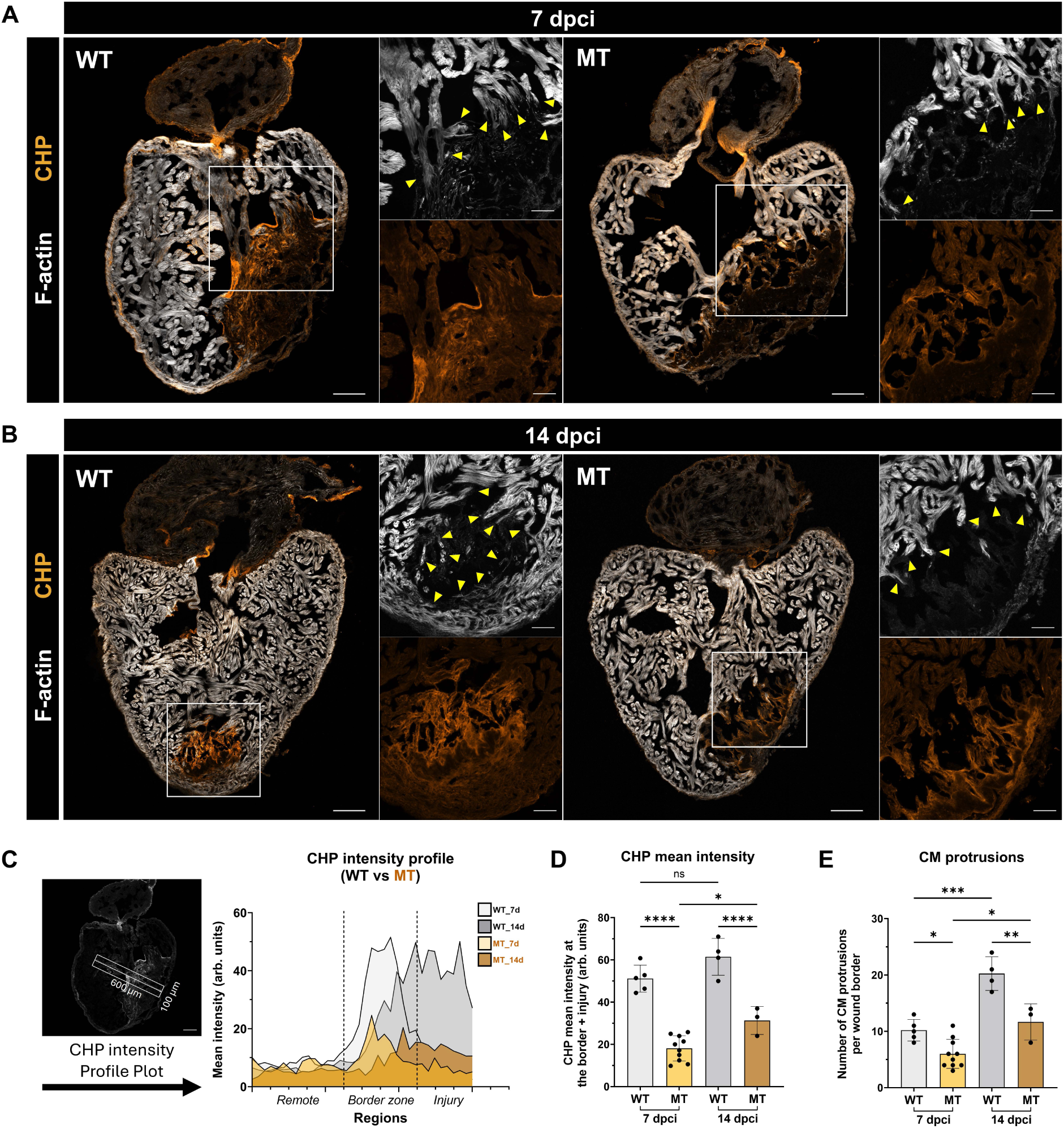
Collagen degradation/remodeling and CM protrusions during heart regeneration. **A-B** Representative images of collagen hybridizing peptide (CHP) staining in WT and *ifng1* MT hearts at **A**, 7 dpci and **B,** 14 dpci, highlighting regions of active collagen degradation/remodeling. Scale bars: 100 µm (overview), 50 µm (insets). **C** Spatial intensity profiles of CHP staining across the remote area, border zone, and injury area at 7 and 14 dpci. Analysis was performed on a 600 µm × 100 µm region perpendicular to the injury axis to ensure consistent sampling of all cardiac zones. **D** Quantification of CHP mean fluorescence intensity within the total injury area at 7 and 14 dpci. **E** Quantification of CM protrusions within the border zone (defined as 200 µm from the injury axis). Data in **C** and **E** are presented as mean ± SD. Statistical significance was determined by two-way ANOVA with Holm-Šídák’s multiple comparison test (\**P* < 0.05, \*\**P* < 0.01, \*\*\**P* < 0.001, \*\*\*\**P* < 0.0001; ns, not significant).

This defect correlated with reduced macrophage accumulation at the injury border zone in *ifng1* MT hearts, despite macrophages being overall more abundant than in WT controls (Supplementary Fig. 3A, quantified in B), and is consistent with the diminished ECM-remodeling gene signatures identified in our macrophage RNA-seq analysis (Fig. 5). Functionally, impaired ECM remodeling in *ifng1* MT hearts was associated with a significant reduction in CM protrusions at the injury border zone (Fig. 6E), critical for CM invasion into and replacement of fibrotic tissue during regeneration^43^. Collectively, these findings provide a mechanistic link between IFN-γ signaling and regenerative ECM remodeling. IFN-γ promotes the activation and proper localization of macrophages with remodeling capacity, thereby facilitating collagen turnover and creating a permissive microenvironment for cardiomyocyte proliferation and invasion during heart regeneration.

### Selective inhibition of IFN-γ signaling in macrophages phenocopies *ifng1* mutants

Given the potential pleiotropic effects of IFN-γ across multiple cardiac cell types, we next sought to determine whether macrophages serve as the primary cellular effectors of IFN-γ-dependent heart regeneration. To address this question, we generated a transgenic zebrafish line, *Tg(hsp70l:loxP-TagBFP-stop-loxP-DN-crfb6-P2A-TagRFP)*^as603^ (hereafter *hsp70l:DN-crfb6*), to enable inducible- and tissue-specific blockade of IFN-γ signaling through expression of a dominant-negative (DN) IFN-γ receptor, *crfb6* (Cytokine Receptor Family B, Member 6; hereafter DN-Crfb6). This truncated receptor retains the extracellular ligand-binding domain, allowing IFN-γ sequestration while preventing downstream intracellular signal transduction^75,76^. To restrict DN-Crfb6 expression to macrophages, we crossed this line with the macrophage-specific Cre driver, *Tg(mpeg1.1:Cre)*^77^ (Fig. 7A). Adult fish were subjected to daily heat-shock induction following cardiac cryoinjury to selectively activate DN-Crfb6 expression in macrophages, and regenerative outcomes were assessed at 7 dpci (Fig. 7B).

**Fig. 7.**
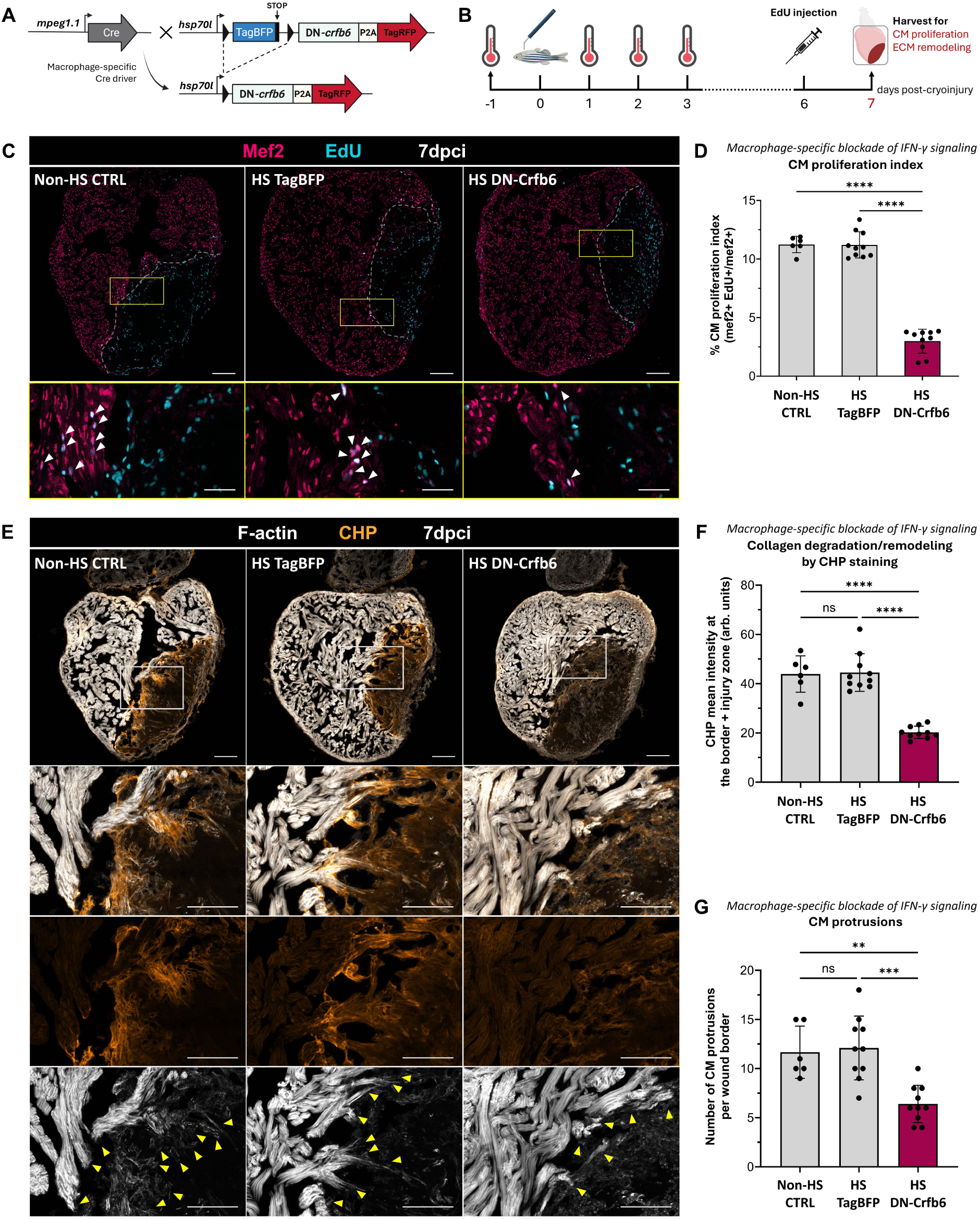
Macrophage-specific blockade of IFN-γ signaling phenocopies global *ifng1* deficiency. **A** Schematic of the Cre-lox transgenic strategy for macrophage-specific inhibition of IFN-γ signaling. The *mpeg1.1:Cre* driver restricts heat-shock–induced overexpression of dominant-negative *crfb6* (DN-Crfb6) to macrophages. **B** Experimental timeline for heat-shock (HS) induction followed by cardiac cryoinjury to assess cardiac regeneration. **C** Representative immunofluorescence images of heart sections at 7 dpci showing Mef2+(magenta) and EdU+(cyan) nuclei. Experimental group include: Non-HS CTRL (genetic control), Tg(*mpeg1.1:Cre, hsp70l:DN-crfb6*) without HS; HS TagBFP (control for HS-induced stress), *Tg(hsp70l:DN-crfb6*) with HS; and HS DN-Crfb6 (macrophage-specific blockade), Tg(*mpeg1.1:Cre, hsp70l:DN-crfb6*) with HS. Proliferating CMs are indicated by overlapping Mef2+/EdU+ signals (violet). Scale bars: 100 µm (overview) and 50 µm (insets). **D** Quantification of the CM proliferation index (%) within the border zone at 7 dpci. **E** Representative images of collagen degradation/remodeling visualized by CHP staining (orange) in the indicated groups at 7 dpci. Scale bars: 100 µm. **F** Quantification of CHP mean fluorescence intensity. **G** Quantification of CM protrusions within the border zone. Sample sizes per group: Non-HS CTRL (n=6), HS TagBFP (n=10), and HS DN-Crfb6 (n=10). Data in **D**, **F** and **G** are presented as mean ± SD. Statistical significance was assessed by one-way ANOVA with Tukey’s multiple-comparison test (\*\**P* < 0.01, \*\*\**P* < 0.001, \*\*\*\**P* < 0.0001; ns, not significant).

Macrophage-specific inhibition of IFN-γ signaling phenocopied the key defects observed in *ifng1* mutant hearts. Hearts expressing DN-Crfb6 in macrophages exhibited a significant reduction in CM proliferation (Fig. 7C, quantified in D) and impaired extracellular matrix remodeling, as indicated by reduced collagen degradation detected by CHP staining (Fig. 7E, quantified in F). Moreover, blocking IFN-γ signaling in macrophages resulted in a marked decrease in CM protrusions at the injury border zone compared with non-heat-shocked and non-transgenic controls (Fig. 7E, quantified in G). Together, these findings demonstrate that IFN-γ signaling acts directly on macrophages to regulate their transition from an inflammatory state to a reparative, ECM-remodeling phenotype, thereby establishing a permissive microenvironment that supports cardiomyocyte invasion and effective heart regeneration.

## DISCUSSION

Successful heart regeneration requires not only CM proliferation but also precise coordination of immune activation, inflammatory resolution, and ECM remodeling^5,32,35,78^. Although immune responses have long been associated with regenerative outcomes across species^12,14,15,38,79,80^, how specific immune signals dynamically instruct macrophage function to enable structural regeneration has remained incompletely understood. Here, we identify IFN-γ as a critical immune regulator that programs macrophage plasticity to enable heart regeneration. By integrating temporal transcriptomics, functional assays, and macrophage-specific genetic blockade, we define a macrophage-intrinsic IFN-γ signaling axis that links inflammatory resolution to regenerative ECM remodeling and CM replenishment.

### IFN-γ as a spatiotemporal regulator of regenerative immune responses

IFN-γ is classically regarded as a potent pro-inflammatory cytokine central to antimicrobial defense and the orchestration of pathological inflammation^24,25,81–83^. In cardiac disease settings, elevated IFN-γ levels have been associated with both adverse remodeling and protective immune activation, leading to seemingly contradictory conclusions regarding its role in repair^25,27,82–88^. Our findings reconcile this paradox by demonstrating that IFN-γ acts in a context- and time-dependent manner during regeneration.

In zebrafish hearts, IFN-γ signaling is dynamically regulated following injury, with early induction of the ligand *ifng1* and subsequent upregulation of receptor components, suggesting a staged signaling program rather than sustained inflammatory activation. Importantly, loss of IFN-γ did not abolish the initial immune response to injury. Instead, *ifng1* mutant hearts mounted an early transcriptional response that largely overlapped with wild-type hearts but failed to sustain reparative programs. This premature termination of the reparative trajectory, accompanied by persistent inflammatory signatures and defective ECM remodeling, indicates that IFN-γ may not be required to initiate inflammation per se, but rather to guide its resolution and productive transition toward regeneration. We further observed a later phase of *ifng1* expression at 14 dpci, coinciding with tissue remodeling and supporting the concept that IFN-γ contributes to macrophage functional switch beyond the acute inflammatory phase. Thus, IFN-γ functions as a dynamic regulator that coordinates immune activation with downstream regenerative events.

### Macrophage reprogramming as the central cellular mechanism downstream of IFN-γ

Our data position macrophages as the primary cellular effectors of IFN-γ–dependent regeneration. Transcriptomic profiling of cardiac macrophages revealed that IFN-γ deficiency constrains macrophages in a pro-inflammatory state characterized by sustained expression of cytokines, antigen-presentation machinery, and complement components. In contrast, wild-type macrophages undergo a marked phenotypic transition toward ECM remodeling and pro-regenerative signaling, including induction of matrix metalloproteinases, collagens, and mitogenic factors known to stimulate CM proliferation. Functionally, impaired macrophage reprogramming in *ifng1* mutants manifests as defective debris clearance, prolonged neutrophil retention, reduced collagen denaturation, and diminished CM protrusion at the injury border zone. These defects converge on a failure to replace fibrotic tissue with regenerating myocardium. Crucially, macrophage-specific blockade of IFN-γ signaling phenocopied the global mutant defects, strongly suggesting that IFN-γ acts through macrophages to control regenerative outcomes. This causal demonstration extends prior correlative studies and establishes macrophage-intrinsic IFN-γ signaling as a key regulatory node in cardiac regeneration.

### IFN-γ links immune resolution to ECM remodeling and cardiomyocyte invasion

A defining conceptual advance of this work is the linkage between cytokine signaling and the structural properties of the regenerative microenvironment. Successful zebrafish heart regeneration requires dynamic ECM remodeling, transitioning from fibrin deposition to collagen accumulation, followed by macrophage-dependent collagen denaturation that permits CM invasion and scar resolution^9,10,43,59,60^.

We demonstrate that IFN-γ–responsive macrophages are required for this spatially and temporally coordinated ECM remodeling program. In wild-type hearts, collagen degradation initiates at the injury border zone and progressively expands across the wound, creating a permissive niche for CM protrusion and tissue replacement. In contrast, *ifng1* mutant hearts exhibit reduced collagen remodeling and diminished CM invasion, resulting in persistence of a collagen-rich scar. These findings place IFN-γ upstream of macrophage-mediated ECM remodeling and support a broader principle: successful regeneration requires not immune suppression, but precise immune reprogramming. IFN-γ ensures that inflammatory activation transitions into a reparative phase that actively remodels the tissue scaffold to enable structural regeneration.

### Evolutionary conservation and translational implications

Our findings build on comparative analyses between regenerative zebrafish and non-regenerative medaka, in which differential immune activation and macrophage behavior were identified as major determinants of regenerative capacity^12,17^ (Chowdhury et al., 2026, *manuscript in preparation*). IFN-γ emerged from these studies as a candidate upstream regulator, including in poly I:C-induced regenerative responses. While poly I:C acts as an exogenous immunostimulant, IFN-γ represents an intrinsic signaling pathway that coordinates macrophage functional transitions during regeneration.

The requirement for macrophages in cardiac regeneration extends beyond teleosts. In neonatal mice, macrophages are essential for regenerative repair, and their depletion results in scarring rather than regeneration^15,89^, a finding consistent with their origin and functional differences compared with their adult counterparts^15,89^. Together, these observations suggest that regenerative capacity may depend less on the presence or absence of immune responses, and more on the ability of macrophages to undergo appropriate functional transitions, an ability that becomes progressively constrained in adult mammals.

From a translational perspective, our study highlights IFN-γ signaling as a potential lever to modulate macrophage function toward regenerative repair. Importantly, our findings caution against simplistic interpretations of IFN-γ as uniformly beneficial or deleterious in cardiac disease. Rather, timing, cellular context, and downstream macrophage state are likely to determine outcomes. Therapeutic strategies aimed at re-engaging regenerative immune programs in the adult mammalian heart may therefore require precise, macrophage-targeted modulation of IFN-γ signaling rather than systemic cytokine manipulation.

### Limitations and future directions

Several important questions remain. First, the downstream signaling pathways and transcriptional networks through which IFN-γ reprograms macrophages toward ECM remodeling and pro-regenerative states warrant further investigation. Second, whether IFN-γ can be harnessed to reprogram mammalian macrophages toward regenerative phenotypes in preclinical settings remains an open and clinically relevant challenge. Finally, understanding how IFN-γ signaling interfaces with mechanical cues, metabolic states, and other cytokine networks will be essential for translating these insights into therapeutic strategies.

### Conclusion

In summary, our study identifies IFN-γ as a central immune regulator that orchestrates macrophage reprogramming to enable debris clearance, ECM remodeling, and cardiomyocyte replenishment during zebrafish heart regeneration. By defining a macrophage-intrinsic IFN-γ signaling axis that couples immune resolution to regenerative tissue remodeling, these findings provide mechanistic insight into how immune signals govern regenerative outcomes and offer new perspectives for promoting cardiac regeneration across species.

## METHODS

### Zebrafish husbandry, handling and ethics statement

All zebrafish procedures were conducted in accordance with institutional and national ethical guidelines. Zebrafish were maintained under standard laboratory conditions following approved Institutional Animal Care and Use Committee (IACUC) protocols at Academia Sinica, Taiwan (protocol no. 24-12-2378). All surgical procedures were performed under tricaine methanesulfonate (MS-222) anesthesia (0.025%), and euthanasia was carried out using 0.2% MS-222 to minimize animal suffering.

### Zebrafish lines

The following wild-type and transgenic zebrafish lines were used in this study: wild-type AB strain, *Tg(fli1:EGFP)^y5^, Tg(-5.1myl7:DsRed2-NLS)^f2^, Tg(mpx:EGFP)^uwm1^, Tg(mpeg1.1:mCherryF)^ump2^, TgBAC(lck:EGFP)^vcc4^, Tg(mpeg1.1:Cre)^fh506^*, and *Tg(hsp70l:loxP-TagBFP-stop-loxP-DN-crfb6-P2A-TagRFP)^as603^* (this study).

The *ifng1* mutant*^as602^* zebrafish line was generated using CRISPR/Cas9-mediated genome editing. Two single-guide RNAs (sgRNAs) targeting exon 1 and exon 3 of the *ifng1* locus were designed to induce a large genomic deletion. Cas9 mRNA and sgRNAs were co-injected into one-cell stage embryos. The resulting allele contained a deletion spanning the intervening genomic region, leading to a frameshift and premature stop codon that produced a truncated, non-functional peptide of 39 amino acids. Founder fish carrying the deletion allele were identified by PCR-based genotyping and outcrossed to wild-type AB fish or relevant transgenic line to establish a stable mutant line. Sequences of sgRNA and genotyping primers are listed in Supplementary Table 1.

The *Tg(hsp70l:loxP-TagBFP-loxP-DN-crfb6-P2A-TagRFP)^as603^* transgenic line was generated to conditionally express a dominant-negative form of interferon gamma receptor under heat-shock control. The construct was derived from *hsp70l:loxP-TagBFP-stop-loxP-DTA-t2A-mCherry^90^* (gift from Stainier’s Lab), with the *DTA-t2A-mCherry* cassette replaced with *DN-crfb6* lacking the intracellular signaling domain while retaining the extracellular and transmembrane domains, followed by a self-cleaving P2A peptide and TagRFP reporter. The *DN-crfb6* open reading frame was amplified by cDNA generated from pooled embryos (6 hpf to 7 dpf) using a reverse primer terminating at the transmembrane domain. The amplified fragment was cloned in-frame upstream of the P2A-TagRFP sequence and inserted into the *hsp70l:loxP-TagBFP-stop-loxP* backbone. The final *hsp70l:loxP-TagBFP-stop-loxP-DN-crfb6-P2A-TagRFP* construct (hereafter *hsp70l:DN-crfb6*) was verified by Sanger sequencing prior to transgenesis. Plasmid DNA was microinjected into one-cell stage wild-type AB embryos together with I-*Sce*I meganuclease^91^.

### Cryoinjury of zebrafish hearts and intraperitoneal (IP) injection

Cardiac cryoinjury was performed as previously described^9,10,92^. Adult zebrafish (4-8 months old) were anesthetized with 0.025% MS-222 (Sigma-Aldrich, A5040). A small incision was made to expose the heart, and a liquid nitrogen–cooled metal probe (−196 °C) was applied to the ventricular apex for 25–30 s. Following the procedure, the fish were transferred to a recovery tank with system water (with water heater set at 28°C) and monitored until they resumed normal swimming behavior. Post-recovery, the fish were returned to the system rack. Hearts were then collected at predetermined time points for subsequent analysis. For *in vivo* labeling of proliferating cells, 10 µL of 20 mM EdU (dissolved in PBS) was administered by IP injection 24 h prior to heart collection.

### RNA isolation for whole-ventricle and isolated cardiac macrophage transcriptomics

For whole-ventricle RNA sequencing, heart ventricles were harvested from WT and *ifng1* MT fish under both cryoinjury and uninjured control conditions. Three ventricles were pooled per experimental group. Briefly, ventricles were carefully dissected in RNase-free ice-cold PBS. Blood clots and excess red blood cells were removed by gently rinsing in several changes of ice-cold PBS using microsurgical tweezers. Following dissection, the pooled ventricles were transferred into 1.5 mL microtubes containing 200 µL of QIAzol Lysis Reagent and immediately snap-frozen in liquid nitrogen (−196°C). For homogenization, an additional 500 µL of QIAzol Lysis Reagent was added to the frozen samples, along with 10-15 zirconium oxide homogenization beads (1.0mm). The tissue was homogenized using a Bullet Blender Storm Pro (Next Advance) set at full power to pulse at 10s intervals for 2 min. Total RNA was subsequently purified using the Qiagen RNeasy Mini Kit according to the manufacturer’s instructions.

For isolated cardiac macrophage RNA sequencing, WT and *ifng1* MT fish in the *Tg(mpeg1.1:mCherryF)^ump2^* background were used. To obtain sufficient macrophages for downstream analysis, approximately 30–35 ventricles were pooled per experimental group. Cardiac tissue dissociation and fluorescence-activated cell sorting (FACS) of mCherry-positive macrophages were performed as previously described^17^. Sorted cells were immediately lysed, and total RNA was extracted using TRIzol™ LS Reagent (Invitrogen) according to the manufacturer’s instructions, followed by library preparation for RNA sequencing.

RNA concentration and purity were measured using a NanoDrop spectrophotometer, and RNA integrity was assessed using an Agilent 2100 Bioanalyzer (RIN scores > 8.0 were used for sequencing). Library preparation and sequencing were performed by local commercial genomic company (Genomics BioSci & Tech Co., Ltd). Briefly, sequencing libraries were prepared using the TruSeq Stranded mRNA Library Prep Kit (Illumina, San Diego, USA) according to the manufacturer’s protocol. Briefly, 1 µg of total RNA was used as input. Validated libraries were pooled and sequenced on an Illumina NovaSeq 6000 platform to generate 150 bp paired-end reads, targeting a depth of approximately 25M reads per sample.

### Whole-ventricle and isolated cardiac Macrophages transcriptomic data processing and analysis

After read mapping, raw read count matrices generated from zebrafish (*Danio rerio*) heart ventricle samples were uploaded to iDEP (integrated Differential Expression and Pathway analysis), a web-based platform for comprehensive RNA-seq data analysis (iDEP v0.9x; http://bioinformatics.sdstate.edu/idep/)^93^, for data pre-processing. Two independent datasets were analyzed: (i) Whole-ventricle RNA sequencing from WT and *ifng1* MT zebrafish collected at uninjured (Un) and post-injury time points (6 h, 1 d, 3 d, 5 d, and 7 d), and (ii) RNA-seq from FACS-sorted cardiac macrophages isolated from WT and *ifng1* MT ventricles at 3 and 7 days post-injury.

Genes with low expression were filtered using a minimum counts-per-million (CPM) threshold of 0.5 in at least one library. Filtered count data were normalized using the DESeq2 median-of-ratios method, and differential gene expression analysis was performed with DESeq2, modeling genotype (WT vs MT) and time point as experimental variables where appropriate. For data visualization and exploratory analyses, normalized counts were transformed using edgeR with a log2(CPM + 4) transformation. Statistical significance was assessed using the Benjamini–Hochberg false discovery rate (FDR) correction, with an adjusted *P* value < 0.05 considered significant. Sample relationships were evaluated by principal component analysis (PCA) and hierarchical clustering, the latter performed using 1 − Pearson correlation with complete linkage, which emphasizes tight co-expression patterns. Functional enrichment analyses, including Gene Ontology (GO) and KEGG pathway analyses, were conducted using the clusterProfiler package^94^ and ShinyGO web-based application^95^ to characterize genotype- and time-dependent transcriptional programs in whole heart tissue and macrophage populations.

### RT-qPCR analysis

Zebrafish hearts were cryoinjured and collected at respective timepoints. Briefly, ventricles from each group were carefully dissected as described above and transferred into 1.5 mL microtubes containing 150 µL of TRIzol Lysis Reagent (Life Technologies Invitrogen, CA). and immediately snap-frozen in liquid nitrogen (−196°C). For homogenization, an additional 850 µL of TRIzol Lysis Reagent Reagent was added to the frozen samples, along with 5-8 zirconium oxide homogenization beads (1.0mm). The tissue was homogenized using a Bullet Blender Storm Pro (Next Advance) set at full power to pulse at 10s intervals for 2 min. Total RNA from homogenate was extracted according to the manufacturer’s instruction. First-strand cDNA was synthesized using Xpert cDNA Synthesis Supermix (GK86.0100, GRISP) according to the manufacturer’s instructions. The RT-qPCR analysis was carried out using PowerUp™ SYBR™ Green Master Mix (A25918, Thermo Fisher Scientific) on a Applied Biosystems™ QuantStudio™ 5 Real-Time PCR System (Thermo Fisher Scientific). The relative gene expression was normalized using *ef1a* as an internal control and fold changes were calculated using the 2^−ΔΔCt^ method. Oligonucleotide sequences for RT-qPCR analysis are listed in Supplementary Table 1.

### Histological analysis, immunofluorescence, and imaging

Adult zebrafish hearts were dissected and fixed in 4% paraformaldehyde (PFA) in PBS for 1 h at room temperature (RT) for AFOG staining and immunofluorescence, or overnight at 4 °C for *in situ* hybridization. Fixed tissues were washed three times in PBS and cryoprotected in a sucrose gradient (15–30% sucrose in PBS) overnight at 4 °C. Hearts were then embedded in O.C.T. (Cryomatrix, Epredia) using dry ice and stored at −80 °C until sectioning. Cryosections (12 µm thickness unless otherwise indicated) were collected using a Leica CM3050S cryostat onto CREST Cre-11 slides (Matsunami, Japan) and stored at −20 °C until use.

For immunofluorescence, cryosections were thawed at RT for 10 min, and O.C.T. was removed by washing in PBS. Sections were incubated in blocking solution (PBS containing 2% goat serum, 0.2% Triton X-100, and 1% DMSO) for 1 h at RT, followed by incubation with primary antibodies diluted in blocking solution overnight at 4 °C. After washing three times with PBSTx (PBS with 0.1% Triton X-100; 10 min each), sections were incubated with fluorophore-conjugated secondary antibodies for 3 h at RT. Nuclei were counterstained with Hoechst 33342 (H3570, Thermo Scientific) at 1:10,000 dilution for 15 min at RT followed by 5 min wash with PBS before mounting with Mowiol® 4-88 (Sigma-Aldrich). For anti-Mef2 staining, antigen retrieval was performed prior to blocking by boiling slides in sodium citrate buffer (10 mM sodium citrate, 0.1% Triton X-100, pH 6.0) at 95 °C for 15 min, followed by gradual cooling to RT. EdU staining was performed using the Click-iT™ EdU Cell Proliferation Kit for Imaging (Alexa Fluor™ 647 dye, C10340, Invitrogen) following the manufacturer’s instructions. Imaging was performed using a Zeiss LSM880 confocal microscope with a Plan-Apochromat 20x/0.8 or Plan-Apochromat 40x/1.3 Oil DIC UV-VIS-IR objectives.

### AFOG staining and scar quantification

Acid Fuchsin Orange G (AFOG) staining was performed to visualize collagen (blue), fibrin (red), and myocardium (orange). Cryosections were thawed at RT for 10 min and post-fixed in Bouin’s fixative for 2 h at 58 °C. Slides were washed in running tap water for 10 min and treated in 1% phosphomolybdic acid (Sigma-Aldrich, HT153) and 0.5% phosphotungstic acid (Sigma-Aldrich, HT152) for 5 min at RT. Sections were then stained with AFOG solution (aniline blue, orange G, and acid fuchsin; Sigma-Aldrich). After staining, slides were dehydrated in 100% Ethanol, cleared, mounted, and imaged. Imaging was performed using a Nikon SMZ25 stereomicroscope equipped with a Nikon DS-Ri2 digital camera. Fibrotic area was quantified by measuring the scar area relative to the total ventricular area from three discontinuous sections per heart.

### Collagen degradation/remodeling and CM protrusions analysis

For collagen hybridizing peptide (CHP) and F-actin staining, 16-µm thickness sections were used. Cryosections were washed twice with PBSTx and blocked in blocking buffer (PBS containing 0.5% Triton X-100, 5% BSA and 1% DMSO) for 30 min at RT. Biotin-conjugated collagen hybridizing peptide (B-CHP; 3Helix) was heat-activated at 80 °C for 5 min prior to dilution and incubated with sections overnight at 4 °C according to the manufacturer’s instructions. After washing, sections were incubated with Alexa Fluor 647–conjugated streptavidin. F-actin was visualized using Alexa Fluor Plus 405–conjugated phalloidin. Quantification of CHP intensity was performed in the total injured area in three nonconsecutive sections exhibiting the largest area from each heart. Quantification of CHP intensity was performed in ImageJ (FIJI) as described previously^43^. CM protrusions were quantified using a maximum intensity projection image from at least 2 sections exhibiting the largest injured area and containing largely trabecular cardiomyocytes. Quantification of protrusions into the injured area were performed in Zeiss ZEN 3.4 (Blue edition).

### Whole-mount *ex vivo* debris clearance assays

Zebrafish ventricles were harvested at the indicated time points and immediately cleaned in ice-cold PBS. Isolated ventricles were incubated in Alexa Fluor Plus 405–conjugated phalloidin (1:500; Invitrogen) for 30 min at 4 °C to label F-actin–associated debris. Following staining, ventricles were washed three times in ice-cold PBS and briefly immersed in 4% PFA to stop cardiac contraction. Whole-mount imaging was performed under Nikon SMZ stereomicroscope.

### Antibodies and reagents

Primary and secondary antibodies, and fluorescent probes used in this study are listed in Supplementary Table 2.

### Quantification and statistical analysis

Data quantification and statistical analyses were performed using Nikon NIS-Elements AR software, ImageJ, and GraphPad Prism (v10). Details regarding quantification methods, statistical tests, and exact *P* values are provided in the corresponding figure legends.

## Data availability

The data supporting the findings of this study are available within the article and its supplemental information. The RNA sequencing datasets generated in this study are available in the NCBI BioProject database (https://www.ncbi.nlm.nih.gov/bioproject) using the access number PRJNA1422991. Further information and requests for resources and reagents should be directed to and will be fulfilled by the lead contact, Shih-Lei (Ben), Lai.

## ACKNOWLEDGEMENTS

We thank the core facilities of the Institute of Biomedical Sciences, Academia Sinica, for technical support, specifically the Light Microscopy, Pathology, and DNA Sequencing facilities (supported by Innovative Instrument Project AS-CFII-113-A12) and the cell sorting services (AS-CFII-114-A9). We are grateful to the members and fish caretakers of the Lai laboratory for their invaluable assistance. We also thank Yao-Ming Chang, Chen-Hui Chen, Li-Chung Hsu, and Kai-Chien Yang for their constructive feedback and discussions.

## FUNDING SUPPORT

This work was supported by Research Project Grants from the National Science and Technology Council (Taiwan) awarded to Ben Shih-Lei Lai (NSTC 114-2320-B-001-010-MY3 and NSTC 113-2923-B-001-003-MY2). Additional support was provided by Clinical Research Collaboration Grant from the Institute of Biomedical Sciences, Academia Sinica (IBMS-CRC108-P01, IBMS-CRC111-P01), the Career Development Award (AS-CDA-112-L04), and the Grand Challenge Project (AS-GC-110-05) from Academia Sinica. Khai Lone Lim was supported by a PhD Fellowship from the Academia Sinica Taiwan International Graduate Program (AS-TIGP) and a PhD scholarship from National Yang Ming Chiao Tung University (NYCU).

## AUTHOR CONTRIBUTIONS

KLL and BSL conceptualized the study and developed the methodology. KLL performed the investigations. YJH performed the bioinformatic analyses. KLL and BSL wrote the original draft and led the editing process. KLL, KC, YJH, and BSL reviewed and edited the manuscript with input from all authors. BSL supervised the study, administered the project, and acquired funding.

## COMPETING INTERESTS

The authors declare no competing interests.

## Notes

### Competing Interest Statement

The authors have declared no competing interest.

