## Supplemental information for "IFN-γ-Dependent Macrophage Reprogramming Coordinates Inflammatory Resolution and Matrix Remodeling in Heart Regeneration"

### Supplementary Information Contents

**Supplementary Fig. 1:** In silico analysis of *ifng1* activation during heart regeneration.

**Supplementary Fig. 2:** Mutagenesis and gross morphology of *ifng1* mutant zebrafish.

**Supplementary Fig. 3:** Spatial distribution of macrophages in WT vs. *ifng1* MT hearts following cardiac injury.

**Supplementary Table 1:** Genotyping PCR, RT-qPCR, ISH and cDNA primer sequences

**Supplementary Table 2:** Primary, secondary antibodies and fluorescent probes used in this study

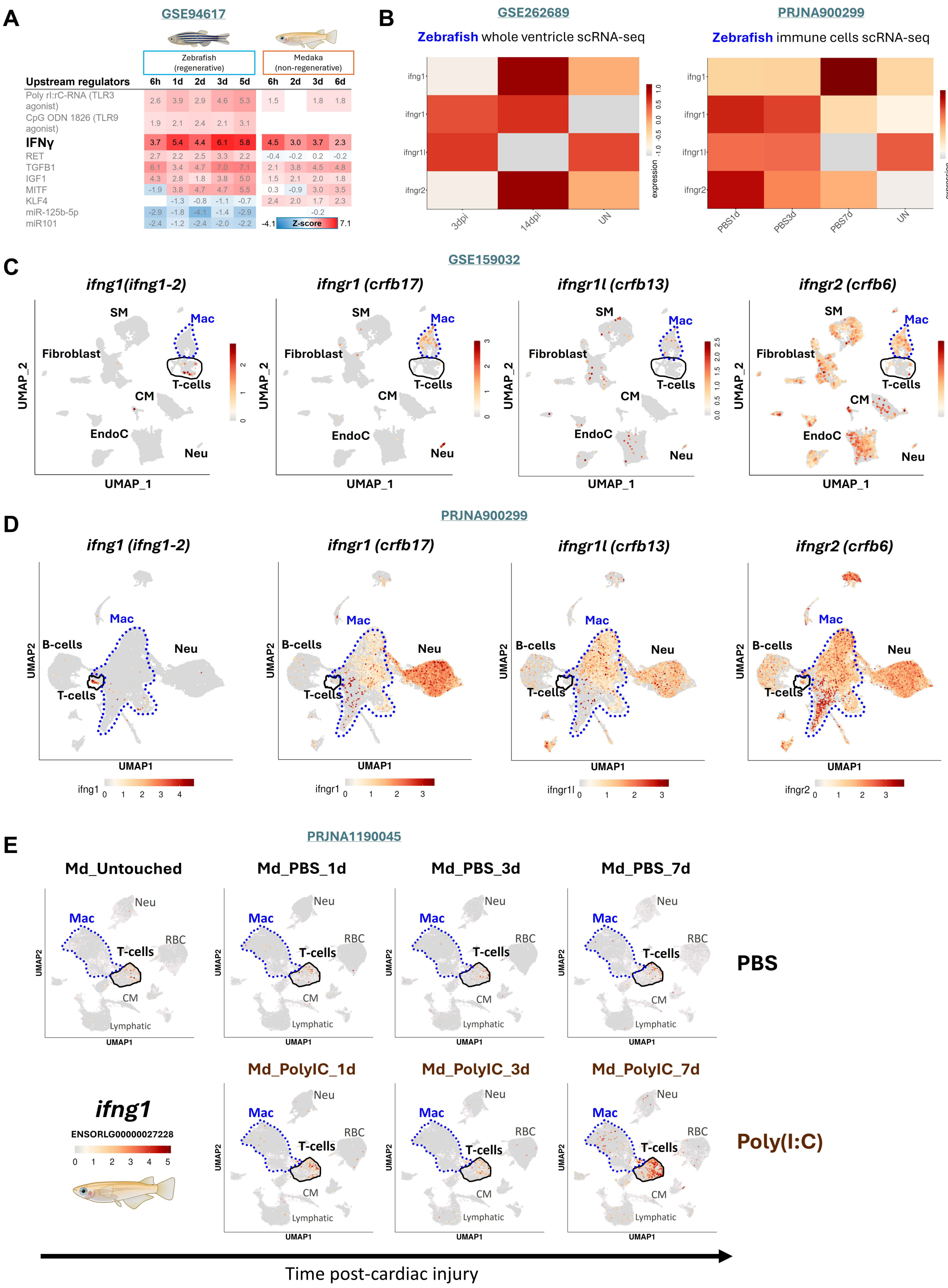

A
GSE94617
Zebrafish (regenerative)
Medaka (non-regenerative)
Upstream regulators
6h 1d 2d 3d 5d 6h 2d 3d 6d
Poly rI:rC-RNA (TLR3 agonist)
CpG ODN 1826 (TLR9 agonist)
IFNγ
RET
TGFB1
IGF1
MITF
KLF4
miR-125b-5p
miR101
Z-score
B
GSE262689
Zebrafish whole ventricle scRNA-seq
PRJNA900299
Zebrafish immune cells scRNA-seq
ifng1
ifngr1
ifngr1l
ifngr2
3dpi
14dpi
UN
PBS1d
PBS3d
PBS7d
expression
C
GSE159032
ifng1(ifng1-2)
ifngr1 (crfb17)
ifngr1l (crfb13)
ifngr2 (crfb6)
SM
Mac
Fibroblast
T-cells
CM
EndoC
Neu
UMAP_1
UMAP_2
D
PRJNA900299
ifng1 (ifng1-2)
B-cells
UMAP1
UMAP2
E
PRJNA1190045
Md_Untouched
Md_PBS_1d
Md_PBS_3d
Md_PBS_7d
RBC
Lymphatic
PBS
Md_PolyIC_1d
Md_PolyIC_3d
Md_PolyIC_7d
ifng1
ENSORLG00000027228
Poly(I:C)
Time post-cardiac injury

**Supplementary Fig. 1 | *In silico* analysis of *ifng1* activation during heart regeneration. A** Ingenuity Pathway Analysis of previously published datasets predicts IFN-γ as an upstream regulator during cardiac regeneration. **B** Meta-analysis of zebrafish single-cell RNA-sequencing (scRNA-seq) datasets reveals reactivation of IFN-γ ligand and receptor expression following cardiac injury. Datasets analyzed include whole-ventricle zebrafish scRNA-seq (GSE262689) and zebrafish immune cells (PRJNA900299). **C-E** Single-cell landscape of: **C**, zebrafish whole-ventricle scRNA-seq (GSE159032); **D**, zebrafish immune cells scRNA-seq (PRJNA900299); and **E**, medaka immune cells scRNA-seq (PRJNA1190045). Feature plots highlight *ifng1* enrichment in T-cell clusters and a subset of macrophages. Expression analysis of IFN-γ ligands and receptors in non-regenerative medaka hearts following poly(I:C) stimulation demonstrates conserved immune responsiveness across species.

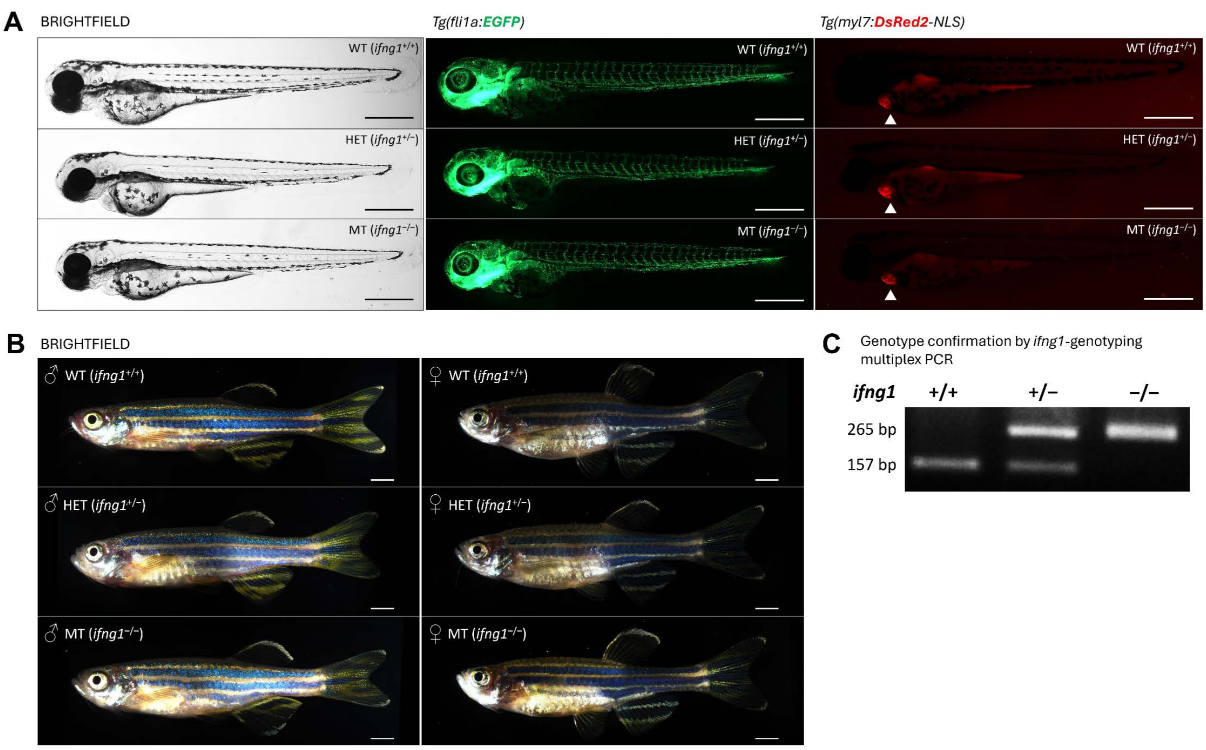


**Supplementary Fig. 2 | Mutagenesis and gross morphology of *ifng1* mutant zebrafish. A** Representative brightfield and fluorescence images of WT, HET and MT larvae at 3 days post-fertilization (3 dpf). No overt developmental abnormalities or deviations in body plan were observed across genotypes. Scale bar: 500 μm. **B** Representative gross morphology of adult of WT, HET and MT fish at 6 months post-fertilization (6 mpf). Scale bar: 2000 μm. Both male and female mutants develop into adulthood without obvious differences in body size or secondary sexual characteristics compared to WT siblings. **C** Representative gel electrophoresis of multiplex PCR genotyping for WT (157 bp), HET (157 bp and 265 bp), and MT (265 bp) alleles.

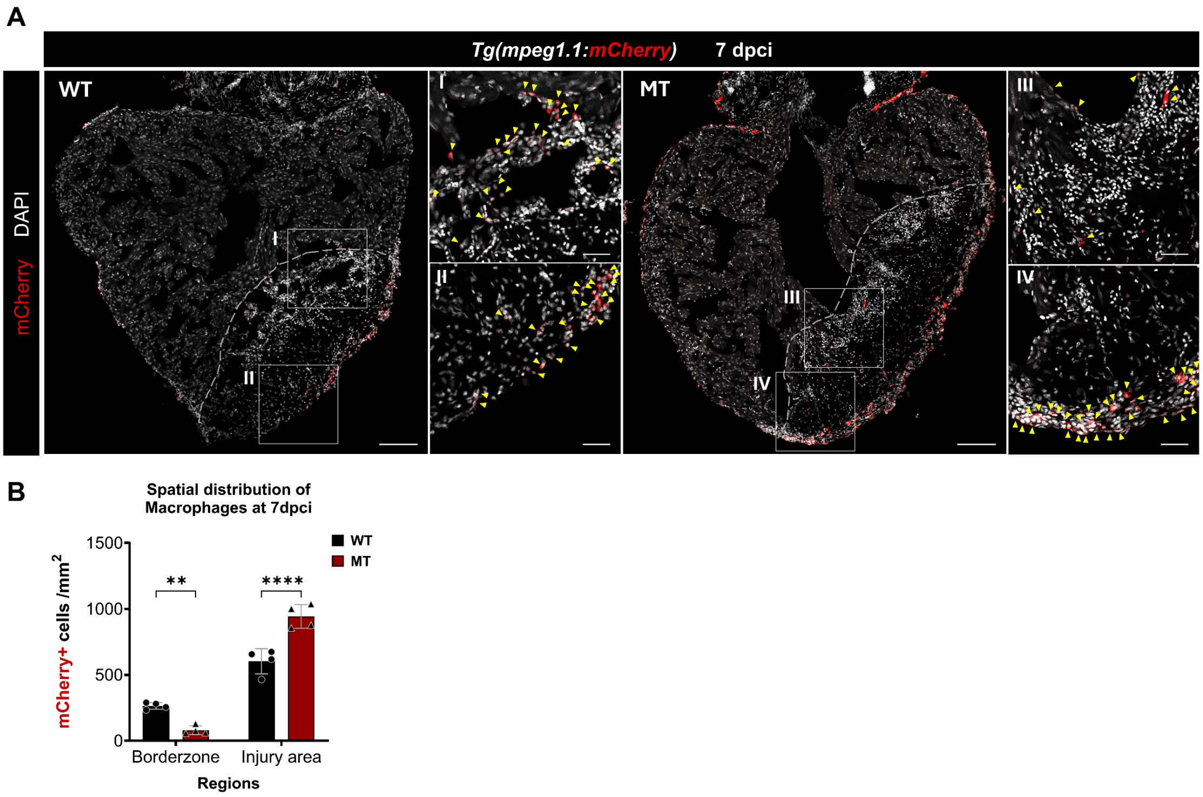


**Supplementary Fig. 3 | Spatial distribution of macrophages in WT vs. *ifng1* MT hearts following cardiac injury. A** Representative immunofluorescence images showing the localization of mCherry+ macrophages in WT and MT hearts at 7 dpci. Insets provide high-magnification views of the border zone (BZ) and injury area (IA), showing individual macrophage density. Scale bars: 100 μm (overview) and 25 μm (insets). **B** Quantification of mCherry+ cell density, normalized to either 200 μm radius of the border zone area or the total injury area (excluding the BZ); WT (n=4) and MT (n=4). Data are presented as mean ± SD. Statistical significance was determined by two-way ANOVA followed by Holm-Šídák's multiple comparison test (***P* < 0.01, *****P* < 0.0001).

**Supplementary Table 1 | Genotyping PCR, RT-qPCR, ISH and cDNA primer sequences**

| **Name** | **Target** | **Sequences (5'-3')** |
|---|---|---|
| *ifng1*-geno-WT_F | Genotyping WT forward | TGGCTGGATTTCCATCGTACGC |
| *ifng1*-geno-WT_R | Genotyping WT reverse | GCCAGGATTTCCAATCTTGAAGGC |
| *ifng1*-geno-MT_F | Genotyping MT forward | CAGCGTGAAAAGGCAACTTTGGC |
| *ifng1*-geno-MT_R | Genotyping MT reverse | TCTCTCACCTTGATCGCCCATAGC |
| *ifng1*-sgRNA-exon1 | *ifng1* exon 1 | GATGCTCTTGTCTAGGTTCTCGG |
| *ifng1*-sgRNA-exon3 | *ifng1* exon 3 | GGATGAAGCTACAAAGGAGAGG |
| *ifng1*-ISH-SP6_F | *ifng1* 5'UTR | ATTTAGGTGACACTATAGAAATGATTGCGCAACACATGATGGG |
| *ifng1*-ISH-T7_R | *ifng1* exon4 | TAATACGACTCACTATAGGGACCTCTATTTAGACTTTTGC |
| *ifng1*-RTqPCR-F | *ifng1* exon1 | CCTAGACAAGAGCATCGAAGAG |
| *ifng1*-RTqPCR-R | *ifng1* exon2 | TCAGGATTCGCAGGAAGATG |
| *crfb17*-RTqPCR-F | *ifng1* exon7 | ATGCGCTCAAAGTGGAGATAG |
| *crfb17*-RTqPCR-R | *ifng1 3'UTR* | CCTTCCGGTGCTTTCTCTTTA |
| *crfb13*-RTqPCR-F | *ifng1* exon5 | TCGTCGTTCTGTGGTTCATC |
| *crfb13*-RTqPCR-R | *ifng1* exon6 | GAGACTCGGGTTGGGAATAAG |
| *crfb6*-RTqPCR-F | *ifng1* exon2 | CTTGACTCCAAAGCAGAGGATAA |
| *crfb6*-RTqPCR-R | *ifng1* exon3 | TGCACGGACCCGAAATAAA |

**Supplementary Table 2 | Primary and secondary antibodies and fluorescent probes used in this study**

### Primary antibodies

| Target | Host species | Catalog number | Supplier | Dilution | Application |
|---|---|---|---|---|---|
| Mef2 | Rabbit | DZ01398 | Boster Bio | 1:150 | Immunofluorescence |
| mCherry | Chicken | ab205402 | Abcam | 1:200 | Immunofluorescence |
| GFP | Rabbit | A-11122 | Invitrogen | 1:200 | Immunofluorescence |

### Secondary antibodies

| Target | Conjugate | Host species | Catalog number | Supplier | Dilution | Application |
|---|---|---|---|---|---|---|
| Rabbit IgG | Alexa Fluor 488 | Goat | A-11008 | Invitrogen | 1:300 | Immunofluorescence |
| Rabbit IgG | Alexa Fluor 568 | Goat | A-11011 | Invitrogen | 1:300 | Immunofluorescence |
| Chicken IgY | Alexa Fluor 568 | Goat | A-11041 | Invitrogen | 1:300 | Immunofluorescence |

### Fluorescent probes and labeling reagents

| Reagent | Fluorophore / Conjugate | Catalog number | Supplier | Working concentration | Application |
|---|---|---|---|---|---|
| Phalloidin | Alexa Fluor Plus 405 | A30104 | Invitrogen | 1:500 | F-actin staining |
| Collagen hybridizing peptide | Biotin-conjugated (B-CHP) | SKU: BIO60 | 3Helix | 20 μM | Detection of denatured collagen |
| Streptavidin | Alexa Fluor 647 | S21374 | Invitrogen | 1:200 | CHP detection |
| EdU detection kit | Alexa Fluor 647 | C10340 | Invitrogen | As per manufacturer | Cell proliferation assay |
| Hoechst 33342 | - | H3570 | Thermo Scientific | 1:10,000 | Nuclear counterstain |
